# Pegasus, a small extracellular peptide regulating the short-range diffusion of Wingless

**DOI:** 10.1101/807701

**Authors:** Emile G Magny, Jose I Pueyo, Sarah A Bishop, Daniel Aguilar-Hidalgo, Juan Pablo Couso

**Author notes:** Current address and author for correspondence.

## Abstract

Small Open Reading Frames (smORFs) coding for peptides of less than 100 amino-acids are emerging as a fundamental and pervasive gene class, found in the hundreds of thousands in metazoan genomes. Even though some of these genes are annotated, their function, if any, remains unknown. Here we characterize the function of a smORF encoding a short 80 aa peptide, *pegasus* (*peg*), which facilitates Wg diffusion during the development of the *Drosophila* wing imaginal disc. During wing development, Wg has sequential functions, and in the later stages, when *peg* is strongly expressed, it patterns the presumptive wing margin. A reduction in Wg protein secretion at this stage produces effects ranging from total abolition of the wing margin to partial loss of bristles and reduction of proneural gene expression. Here we show that the Peg peptide enhances the short-range of Wg diffusion in this context, in order to produce a proper wing margin. We show that CRISPR/Cas9-mediated mutations of *pegasus* generate wing margin phenotypes, and changes in target gene expression, consistent with a role in Wg signalling. We find that Peg is secreted, and that it co-localizes and co-immunoprecipitates with Wg, suggesting that Peg directly binds Wg in order to enhance its signalling, and our data from fixed and *in-vivo* Wg gradient measurements supports a model in which this enhancement occurs by increasing diffusion of extracellular Wg. Our results unveil a new element in the regulation of the Wg signalling pathway, and shed light on the functional consequences of the miss-regulation of Wg diffusion in this developmental context, while also reminding us of the functional diversity, and relevance of small open reading frame genes.

## Introduction

Small Open Reading Frames (smORFs) coding for peptides of less than 100 amino-acids are emerging as a fundamental and pervasive gene class, mostly uncharacterised but found in the tens of thousands in metazoan genomes (*1–3*)(*4*). A rich source of smORFs are the so-called long-non-coding RNAs, which several studies indicate are in fact translated in many cases (*5–7*), and which have already been shown to produce peptides with important functions (*8–12*). There is also a small number of genes annotated as coding for small peptides (short CDSs) whose function is also unknown. Their short peptides are about 80 aa long with a propensity to include both positively charged domains and hydrophobic transmembrane α-helixes (TMHS), typical of secreted and membrane-associated peptides (*2, 5*). Characterised shortCDS peptides localise at or near membrane organelles, and their known functions range from antimicrobial peptides (*13*), organelle components (*14–16*), and cell signals (*17–19*). We identified a number of apparently translated shortCDSs in *Drosophila* for full functional characterisation, including CG17278, which we call *pegasus* (*peg*). Here, we show that *peg* encodes a secreted peptide that binds and modulates the cell signalling Wnt protein Wingless.

The development of the fly wing is a well-studied developmental system, that led to the characterisation of several important cell signals in animal development. Amongst these signals is the secreted *wingless/Dwnt1* protein (Wg). The Wg protein has sequential expression patterns and functions in fly wing development (*20*), and at the end of larval development it patterns the presumptive wing margin. Thus, reducing Wg protein secretion or transport with Wg mutant proteins produces effects ranging from total abolition of the wing margin to partial loss of bristles and reduction of proneural gene expression (*21, 22*). Here, we show that the Peg peptide increases the short-range of Wg signalling by increasing Wg diffusion, and that this range enhancement is essential to establish the full extent of proneural gene activation, and hence, the full wing margin pattern.

## Results

### 1-Identification of *peg*

*peg* encodes an 80 aa peptide with a signal peptide domain and a Kazal2/Follistatin-like domain (FS-like), which localised to vesicular-like cellular structures in S2 cells. FS-like domains are present in Agrin and other large heparin-sulphate proteoglycans (HSPGs). HSPGs are secreted extracellularly and bind to the extracellular matrix, facilitating the anchoring or diffusion of other proteins across it; in particular, HSPGs have been shown to promote Wnt and Hh signalling in vertebrates and flies (*23*). *peg* showed good conservation in insects, and possible conservation in vertebrate genes related to the SPINK extracellular protease inhibitors (Fig. 1A). *peg* was strongly expressed in the developing fly wing but was excluded from the presumptive wing margin (Fig. 1B-C), a pattern complementary to that of the fly Wnt gene *wingless* (*wg*) (Fig. 1B-C) (*21, 24*).

**Figure 1.**
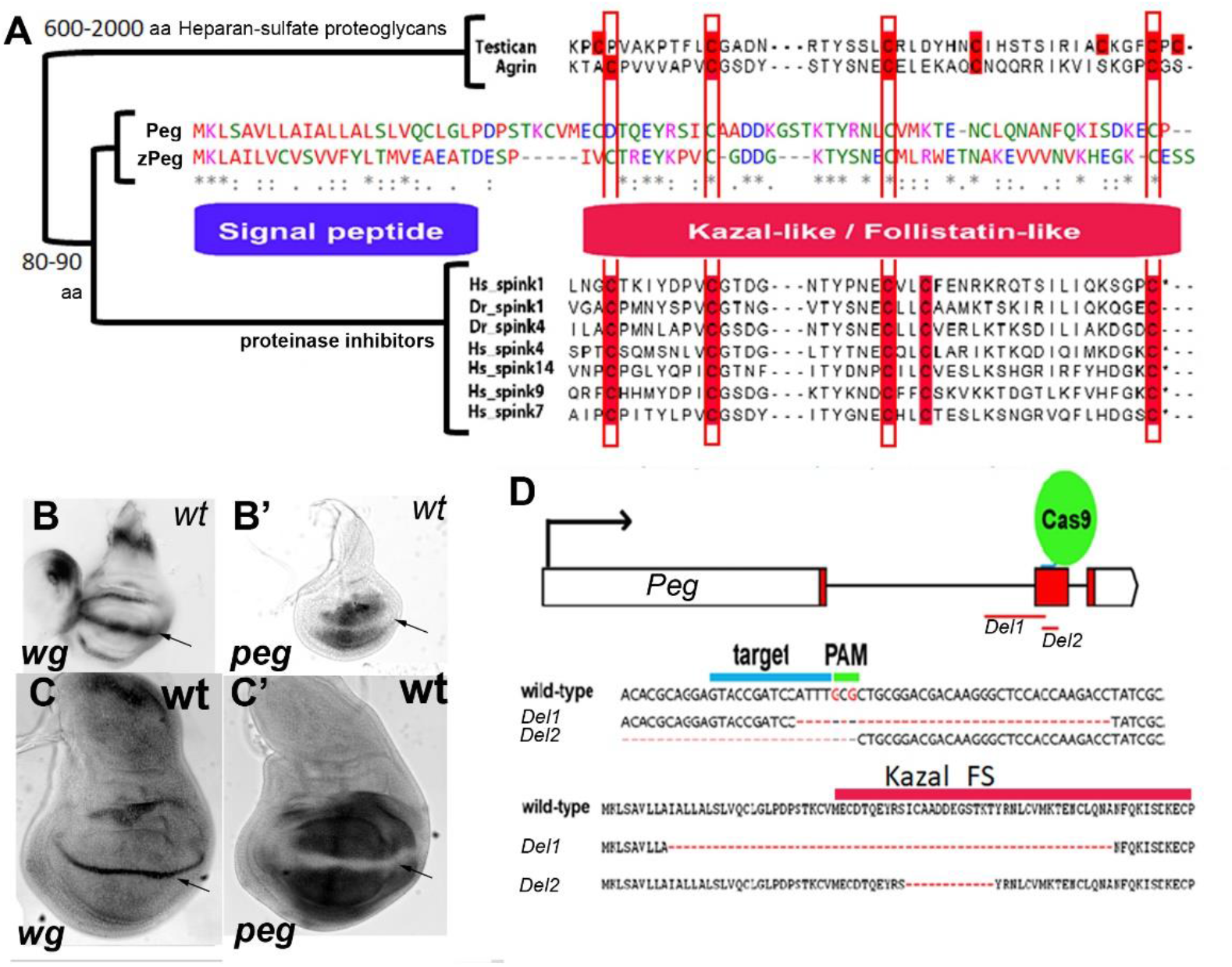
**A.** Phylogenetic tree and alignments of different proteins with FS-like domains (including Peg, the large secreted proteoglycans Agrin and Testican, and the family of small secreted Serine Protease Inhibitors of the Kazal type (Spinks)). Note the closer relationship between Peg and zPeg, a putative Peg homologue in zebrafish. Note also that the Fs-like domain of the protease-inhibitory Spink family shows a highly conserved pattern of cysteines (highlighted in red), whereas the pattern in Peg resembles more that of the non-inhibitory proteoglycans. The signal peptide and FS-like domains are indicated in blue and pink, respectively. **B-C.** *In-situ* hybridization showing the patterns of expression of *wg* and *peg* in mid (B, B’) and late (C, C’) third instar wing imaginal discs. *peg* is strongly expressed in the wing pouch, but not in the cells of the presumptive wing margin (arrows), where Wg is expressed at these stages. **D.** Diagram showing the genomic site targeted for CRISPR/Cas9 mutagenesis, the DNA and protein sequence of the generated null alleles Del1 and 2, and the Kazal domain.

### 2-Characterisation of the *peg* mutant phenotype

We generated null mutants by CRISPR-Cas9 producing small deletions within the coding sequence of the peptide (Fig. 1D). These mutants behave as genetic nulls and show high pupal lethality and a significant reduction in the number of chemosensory bristles at the wing margin (Fig. 2 A,D), a characteristic phenotype of *wg* loss of function (*21*). The sensory organ precursors (SOPs) that give rise to chemosensory bristles are determined at the third instar larval stage of development, from cells within a proneural field induced by high levels of Wg signalling, received by a row of about 3 cells directly neighbouring the row of Wg expressing cells at the presumptive dorso-ventral boundary (*21*). In *peg* null mutants, the proneural marker *senseless* (*sens*) (*25*) showed a significant reduction in the width of the proneural field consistent with the observed reduction of chemosensory bristles phenotype (Fig. 2B, Sup. Fig. 1A,B). Altogether the *peg* phenotype suggests that Peg acts as a positive factor in Wg signalling.

**Figure 2.**
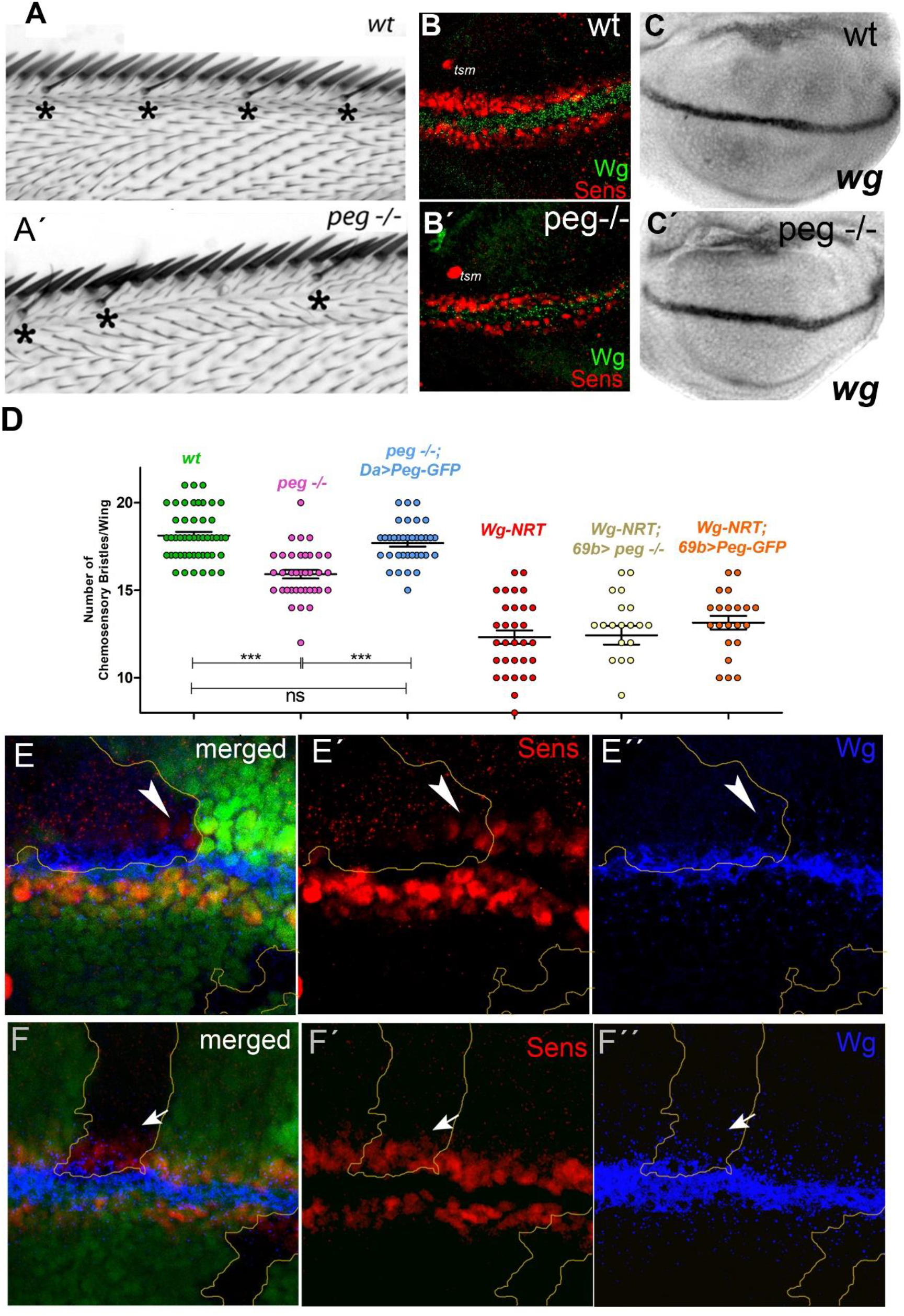
***A.** peg* mutants (A’) present a reduction in wing margin sensory organs (dorsal chemo-sensory bristles (*)) compared to wild type (A). **B.** The expression of *sens* in late third instar wing imaginal discs shows a reduction in *peg* mutant flies (B’) compared to wt (B). Note the the unaffected twin campaniform sensila (tsm) precursor in both images. ***C.** In-situ* hybridization showing that *peg* null mutants (C’) present no changes in Wg mRNA expression compared to wt (C). **D.** Quantification of chemosensory bristles in different genetic backgrounds. *peg* mutants show a significant reduction compared to wild-type. This phenotype is rescued by expression of Peg-GFP. Flies homozygous for a membrane bound version of Wg (Nrt-Wg) show a similar, albeit stronger, phenotype, which is not rescued by over-expression of Peg-GFP. **E-E”.** Large *peg* mutant clones (labelled by absence of GFP, green) show a reduction in *sens* expression (red) and in extracellular Wg (blue dots) present outside the wg-expressing cells (blue intracellular staining, which remains normal in the clone). Within these large clones, mutant cells neighbouring wild-type cells have a near-normal expression of *sens* and extracellular Wg (Arrowheads). **F-F”** Small clones (up to 6 cell diameters wide) show no reduction in either *sens* expression (F,F’) or Wg diffusion (F,F”) (arrows). Note that intracellular Wg is always normal.

*peg* encodes a possible signal peptide domain, suggesting that is a secreted peptide. We looked at the expression of *sens* in *peg*−/− clones, to assess the cell autonomy of Peg, hypothesising that if Peg is indeed secreted it may act in a non-autonomous way. We also assessed the expression of Wg in order to determine if it could be a target of Peg. We observed no effect in the proneural field within small *peg* mutant clones (Fig. 2F), but in larger clones there is a loss of *sens* in cells located more than 3-4 cell diameters away from the nearest Peg-producing cells (Fig. 2E). Interestingly in these large mutant clones we also observe a reduction in extracellular Wg, while in the smaller clones with normal *sens* expression the diffusion of Wg appears normal (Fig. 2F). These results show that Peg indeed acts in a cell non-autonomous way, consistent with its predicted secretion, with a range of some three cells. Moreover, these results show that Peg is required for proper Wg protein allocation. Due to its positive effects on Wg signalling and transport, we called this gene *pegasus* (*peg*), after the mythical winged horse that transported Greek heroes.

### 3-Peg-GFP is functional, secreted and co-localises with Wg

To establish the cellular localisation of Peg and its relationship with Wg *in vivo*, we generated transgenic flies expressing a *UAS-Peg-GFP* fusion (methods). Peg-GFP rescues the chemosensory phenotype of null *peg* without producing unrelated phenotypes, showing that Peg-GFP is a bona-fide replica of native Peg. We expressed Peg-GFP with *ptc-GAL4-Gal80*^*ts*^ in a stripe across the dorso-ventral axis and the Wg expression stripe. We observed Peg-GFP shortly after induction of its expression (5h), and by 12 hours it was detected outside this domain, showing that Peg is indeed secreted from the *ptcGal4 UAS-Peg-GFP* expressing cells (Fig. 3A, and Sup. Fig. 2). By co-expressing *UAS-Peg-GFP* and *UAS-Wg*, we found that Peg co-localises with Wg, both in expressing cells and in neighbouring ones (Fig. 3B, C). Importantly, Peg-GFP also co-localises with endogenous Wg, as corroborated when Peg-GFP is expressed throughout the wing pouch with *rn-Gal4* (Sup. Fig. 2). To test for a direct protein-protein interaction between Peg and Wg we performed a co-immunoprecipitation from protein extracts of third instar larvae expressing Peg-GFP. Endogenous Wg was retained by Peg-GFP, but not by GFP-only controls, showing that Peg can bind directly and specifically to Wg *in vivo* (Fig. 3D).

**Figure 3.**
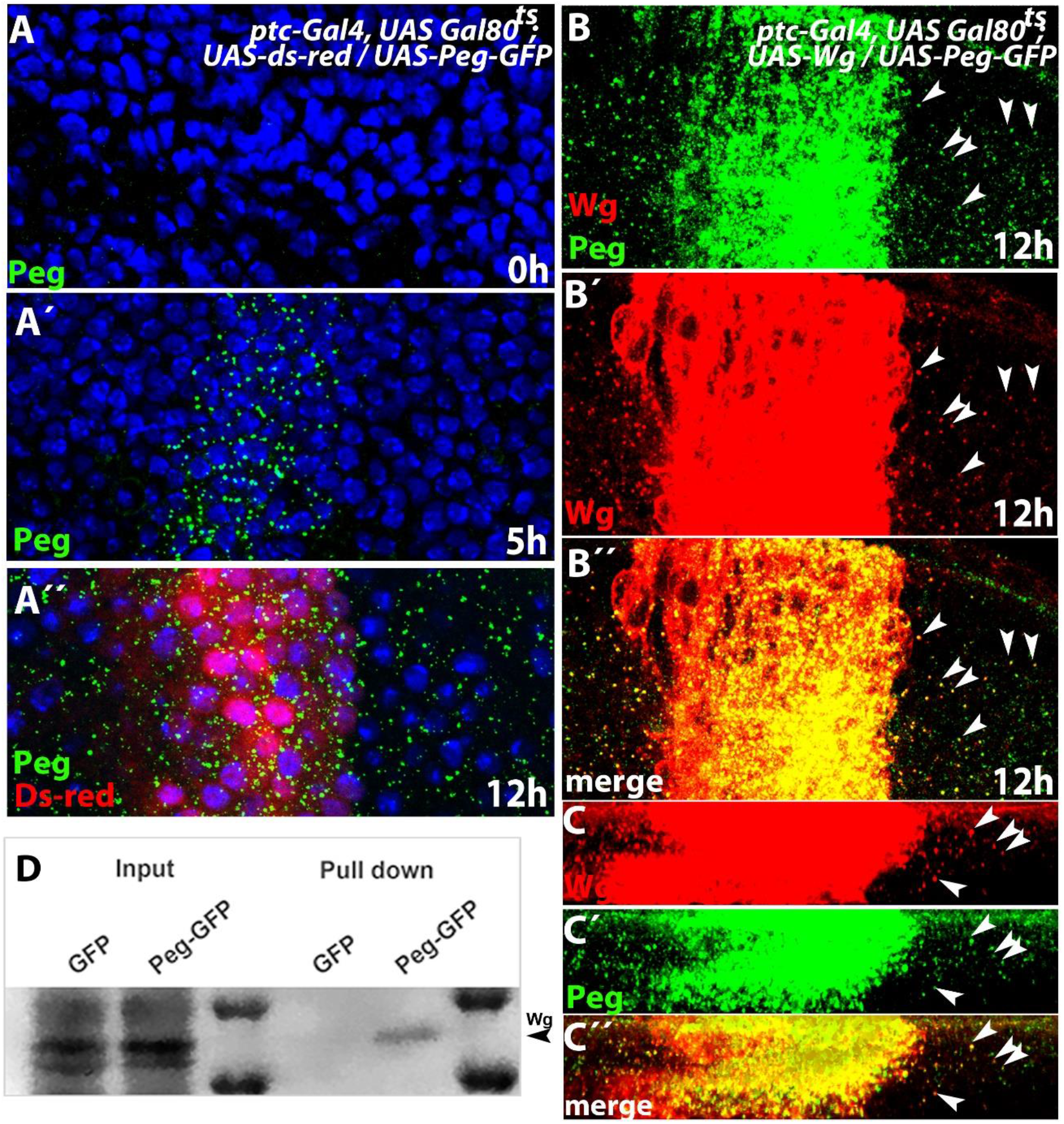
**A.** Induction of Peg-GFP expression with *ptc-Gal4-UAS-Gal80*^*ts*^ by temperature shift (18°C to 29°C) prior to dissection shows that Peg-GFP (green) is secreted. (A) Without temperature shift no GFP expression is detectable; (A’) after a brief 5h shift, GFP signal becomes detectable in the *ptc* expression domain; (A”) 12 hours after temperature shift, secreted Peg-GFP can be detected outside of the *ptc* expression domain (labelled with Ds-red). **B.** When co-induced, Peg-GFP and Wg, are similarly secreted from the *ptc* expression domain, and show extensive co-localisation (arrowheads). Temperature shift 12h prior to dissection. **C.** Orthogonal view of (B), highlighting the co-localisation between Peg-GFP and Wg (arrowheads). **D.** Endogenous Wg co-immunoprecipitates with Peg-GFP, but not with GFP-only controls, showing that a specific direct protein-protein interaction exists between Peg and Wg.

### 4-Peg facilitates Wg transport

Having established that the Peg peptide is secreted, localizes with and binds extracellular Wg, we focused on the effect of Peg on the transport and function of extra-cellular Wg. In the absence of Peg, Wg showed a narrower range of transport compared to wildtype (wt) (Fig. 4A, B, F), consistent with the narrower *sens-*expressing proneural field, lack of chemosensory bristles observed in these mutants, and the lower Wg diffusion observed in large *peg*−/− clones (Fig. 2A-D). Reciprocally, when Peg-GFP was over-expressed in the posterior wing compartment, we observed extended Wg diffusion (Fig. 4C,F’) and Sens expression (Fig. 4D-E’, F”). Peg has a similar effect on Wg-GFP transport and signalling, when UAS-Wg-GFP is either co-expressed with UAS-Peg, or expressed in a *peg* mutant background, using *ptc-Gal4* (Fig. 5A-A”). We used the *ptc-GAL4-Gal80*^*ts*^; *UAS-Wg-GFP* expression system to perform Fluorescence Recovery after Photobleaching (FRAP) and quantify the transporting effect of Peg *in vivo*. We observe in third instar imaginal discs a recovery of 40% of the Wg fluorescence intensity by 5 minutes (Fig. 4 G,H), which suggests very fast Wg transport dynamics, whereas *peg* mutants show a slower rate of Wg recovery, reaching only 14% by the same time (Fig. 4 G’,H).

**Figure 4.**
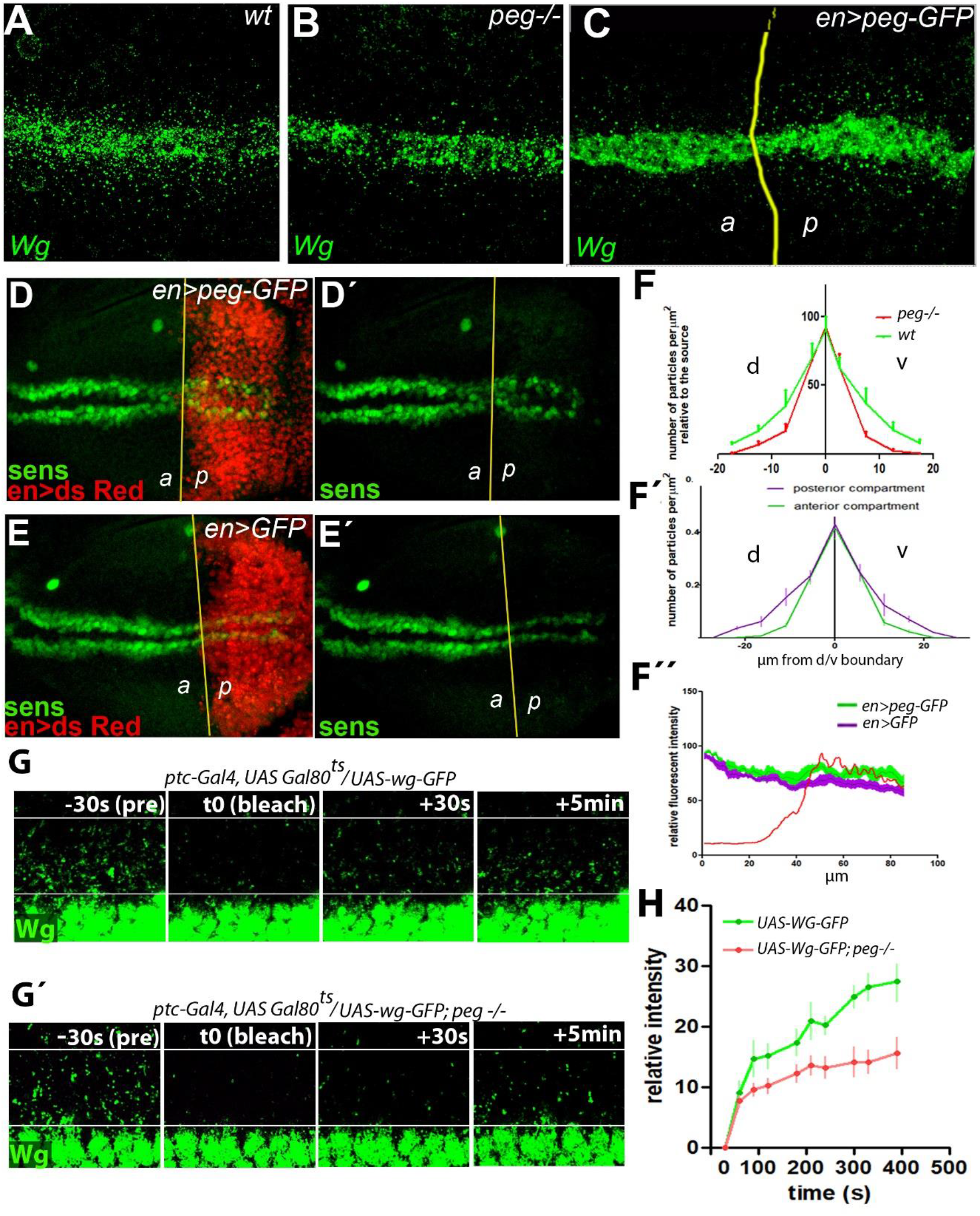
**A-C.** Wg diffusion is reduced in *peg* mutant wing imaginal discs (B) compared to *wt* (A), and it is enhanced in the posterior compartment when *peg-GFP* expression is driven by posterior-specific driver *en-Gal* (C). **D-E’.** Over-expression of Peg-GFP in the posterior compartment (D-D’) leads to a compartment-specific increase in *sens* expression, compared to control imaginal discs over-expressing GFP-only with the same driver (E-E’). tsm, twin campaniform sensila. **F.** Quantification of Wg (F,F’) and Sens (F”) signal intensity from panels A-E”. (F, F’) quantification of the average number of fluorescent Wg particles per μm^2^ on either side of the presumptive wing margin center (line at 0 μm). (F) wing discs of wt flies show significantly higher number of fluorescent particles, further away from the center than those of *peg*−/− mutants. (F’) The posterior compartment of *en-Gal4, UAS-peg-GFP* wing discs show a significantly higher number of fluorescent particles further away from the center than the anterior compartment in the same wing discs. (F”) The posterior compartment of *en-Gal4, UAS-peg-GFP* wing discs show significantly higher levels of Sens signal than the posterior compartment of *en-Gal4, UAS-GFP* controls. **G-G’.** *In-vivo* imaging showing Wg-GFP before (pre), immediately after (t0, bleach), 30s, and 5 min after photo-bleaching (performed on indicated region within white squares), in a wild-type (G) or a *peg* mutant (G’) background. Note the much slower recovery of GFP signal in the *peg* mutant background. Expression was induced 24h prior to dissection. **H.** Quantification of the fluorescence recovery from the experiments shown in panels G-G’, showing the average fluorescence intensity within the bleached region, over time, and relative to the fluorescent intensity before bleaching (pre). Acquisition was made every 30s. t0 indicates the time immediately after bleaching.

**Figure 5.**
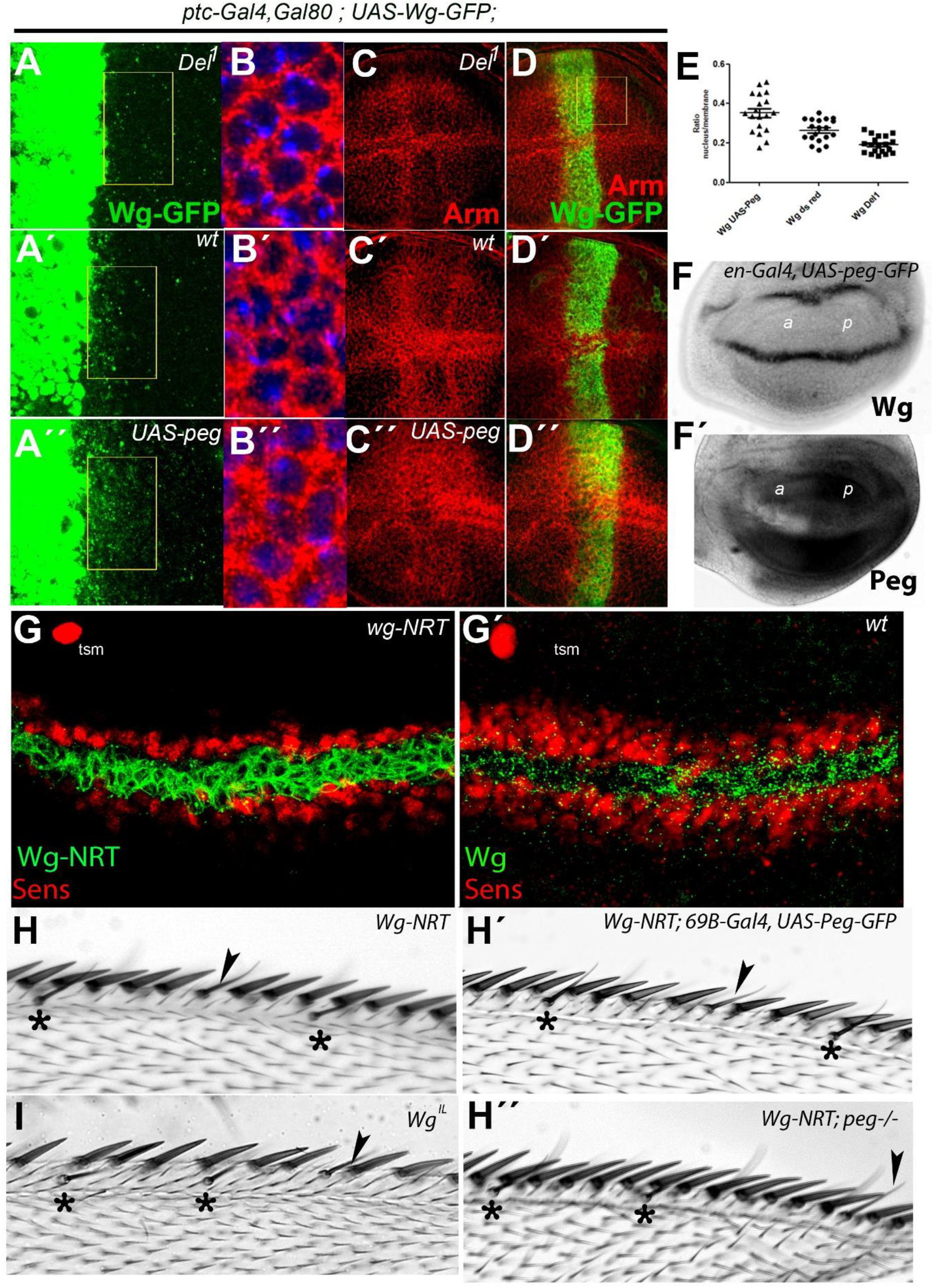
**A-A”.** Induction of Wg-GFP expression with *ptc-Gal4-UAS-Gal80*^*ts*^ by temperature shift 12 h before dissection in a *peg* mutant background (A), a wt background (A’), or in co-induction with Peg *(A”)*. Diffusion is reduced in a *peg* mutant background, and enhanced when Wg is co-expressed with Peg. **B-B”.** High magnification of the cells neighbouring Wg expressing cells (yellow rectangles in (A-A”)), stained with anti-Armadillo (red) and DAPI (blue). Nuclear Armadillo levels are reduced in *peg* mutants (B), and enhanced in wing discs co-expressing Peg *(B”)* compared *to wt (B’)*. **C-D”.** Lower magnification images showing non-membrane accumulation of Armadillo (red) in the cells near Wg-GFP (green) expression in *peg* mutant (C, D), wild-type (C’,D’), and Peg over-expression backgrounds. Accumulation of Arm is lowest in the absence of Peg, and highest when it is overexpressed. **E.** Quantification of the ratio of signal from nuclear/membrane Armadillo from panels B-B”. **F-F’.** *In-situ* hybridization showing *wg* (F) or *peg* (F’) mRNA expression when *peg-GFP* is over-expressed in the posterior compartment. *peg* over-expression is noticeable as the probe hybridizes both endogenous *peg* and induced *peg-GFP* mRNAs, yet it has no effect in Wg mRNA expression. *a* and *p* indicate anterior and posterior compartments. ***G-G’.*** In *wg-NRT* homozygous discs (G) Wg (green) localises to the cell-membrane, and fails to diffuse away from the expressing cells. In these discs there is a strong reduction in *sens* expression (red) compared to wild type (G’). tsm, twin campaniform sensila precursor. ***H-H’.** wg-NRT* homozygous flies (H) have a “narrow double row” phenotype (Couso et al. 1994) presenting a reduction in slender chemosensory bristles (*) and stout mechanosensory receptors (sometimes replaced by remaining chemosensory bristles, arrowhead). Over expression of *peg-GFP* in the wing imaginal disc using the *69B-Gal4* driver (H’), or using a *peg*−/− background (H”) does not affect the *wg-NRT* wing margin phenotype, (See Fig. 2D for quantification). Flies homozygous for the *wg*^*IL*^ temperature sensitive allele, grown at the permissive 17°C temperature, show a similar lack of bristles phenotype (I), interestingly this allele has been shown to affect Wg secretion(*21, 30*).

### 5-Direct effect of Peg on Wg protein function

Our results suggested that Peg acts extracellularly and directly on Wg by enhancing its transport dynamics as required for proper wing margin patterning, including proneural gene activation. Next, we stablished that the positive effect of Peg on Wg transport and signalling are linked and map to the Wg protein. Firstly, we carried out *in-situ* hybridisations to detect *wg* mRNA, and we did not observe an expansion of the *wg* transcription domain upon over-expression of Peg, in the posterior compartment using our *en-Gal4, UAS-peg-GFP* line (Fig. 5F). Secondly, the correlated expansion of Wg expression and of its signalling target *sens* were linked by a similar expansion of Armadillo (Arm) nuclear localisation, a known marker of Wg signalling activity (Fig. 5A-E). Thirdly, abolition of Wg transport was epistatic over Peg function: it was shown that a substitution of endogenous Wg protein with a membrane-tethered version (Wg-NRT) gives rise to apparently morphologically normal flies (*22*). We looked in detail at the Wg margin patterning and *sens* expression in these Wg-NRT flies and found similar phenotypes as those of *peg* mutants (Fig. 5G-H’), which over-expression of Peg was not able to rescue, and which loss of function of *peg* did not enhance (Fig. 5H’-H”).

Overall our results show that Peg promotes Wg transport on the one hand, while on the other hand Peg promotes Wg signalling acting upstream or on the Wg protein itself. The simplest hypothesis is that both effects are causally linked, such that the Peg peptide facilitates Wg protein transport, which in turn results in expanded Wg signalling and expression of target genes. Thus, our results corroborate that a local spatial range of Wg of 3-4 cells is required for proper establishment of the wing margin and its associated proneural field (Couso et al. 1994), and that reducing Wg transport by either Peg loss or Wg protein alterations results in a reduction of these developing structures (Fig. 5H-I).

### 6-Effect of Peg on Wg diffusion

Since Peg has a predicted domain related to both protease inhibition activity (*26, 27*), and HSPGs, we asked whether Peg may have an effect in Wg stability/degradation (working as a protease inhibitor), or in its extracellular movement (working as an HSPG). Using the data from our endogenous Wg profiles in wt and *peg* mutants (Fig. 4F), and our Wg FRAP experiments (Fig. 4G-G’), we formalized the effective transport dynamics of Wg using the reaction diffusion model of Kicheva et al. (*28*)(see methods). We obtained a wild-type decay length for Wg (λ_wt_ = 6.14 ± 1.31 μm) similar to Kicheva’s (5.8 ± 2.04 μm), but a significant reduction in *peg* mutants (λ_peg_= 3.19 ± 0.57 μm), together with a significant reduction in the Wg diffusion coefficients between *peg* and wt: D_wt_ = 0.52 ± 0.08 μm^2^/s and D_peg_ = 0.15 ± 0.06 μm^2^/s. Further, we found no significant difference in the effective degradation rates between wt and *peg* (K_wt_ = 0.014 ± 0.006 /s, K_peg_ = 0.015 ± 0.008 /s). These results imply that Wg diffuses at a slower rate in *peg* mutants, without changes to Wg degradation. To corroborate these calculations, we measured directly the levels of mRNA and protein using qRT-PCR and quantitative Western blots. Consistent with our *in-situ* hybridisation data (Fig. 5F), we observed no significant changes in mRNA transcript levels in *peg* mutants, nor when *peg* was over-expressed (Sup. Fig.2C). Similarly, and consistent with our imaging data, we observed no significant differences in Wg protein levels in the wing pouch of imaginal discs of *peg* mutants, or in those over-expressing UAS-Peg. Altogether our cellular, biochemical and mathematical data support a role for Peg as a facilitator of Wg diffusion.

## Discussion

Although Wg has been characterised as a diffusible ligand, acting on a range of several cells (*29, 30*), it was reported that the substitution of the *wg* locus by a membrane tethered version of Wg (Wg-NRT) gave rise to flies with overtly normal morphology (although poorly viable), suggesting that diffusion of Wg may not be actually required during development (*22, 31*). However, some studies using Wg-NRT have shown that Wg signalling at a distance is required in specific contexts (*32, 33*). Here we show that short-range diffusion of Wg is indeed required for the appropriate patterning of the wing margin, as initially reported (*21*). It has been repeatedly shown how Wg has sequential short range signalling functions in *Drosophila* wing development, (instead of a single, long range morphogenetic function), a crucial fact that reconciles results from several groups (*20, 21, 34, 35*)(*36–39*) (Sup. Fig 3). Thus, early in wing development Wg establishes a 10-cell wide wing field from the ventral compartment; Wg perturbations at this stage result in either loss of the entire wing, or its duplication (*20, 34, 36, 40*). Shortly afterwards, transitory expression of Wg across the entire developing wing is required for wing growth (*20, 37, 41*)(*31*), and finally, Wg is expressed in a stripe along the dorsal-ventral boundary of the wing from where it patterns the wing margin (*21*)(*22*). The widespread growth function does not strictly require Wg diffusion (since Wg is then transcribed in nearly all cells of the developing wing) but the other two functions rely on effective short-range Wg signalling. The early establishment of the wing field occurs in a primordia 10 cells across, yet by the end of development two days later, its effects are inherited and indirectly felt across hundreds of cells; whereas the late patterning of the wing margin offers an immediate and direct read-out of the functional range of Wg signalling. In this late context, the Peg peptide is an essential contributor to the short-range Wg signal. Peg enhances Wg diffusion to reach cells located 3-4 cell diameters from the Wg source, which otherwise would be deprived of the Wg signal. The signalling to the entire proneural field is essential to ensure the development of all the neural precursors and bristles that the wing margin displays, in particular the number of chemoreceptors that inform the fly of its surroundings, and the subsequent positioning of the mechanoreceptors that allow the fly to keep steady constant flight. The role of the Peg small secreted peptide, together with those of canonical proteins, corroborates that transport of the Wnt developmental signals is a complex and highly modulated process (reviewed in (*42*), different from the elegant classical models of freely diffusing morphogens (*43*).

Our results also corroborate the growing importance of smORFs. Given the characterised examples (*12, 17–19, 44*), reviewed in (*4, 45*), and their numbers and molecular characteristics (*2*), it is likely that more smORFs of importance in development and inter-cellular communication await characterisation across metazoans. Only continued smORF studies will uncover the true answers to their potential.

## Materials and methods

### -Drosophila lines

Fly stocks and crosses were cultured at 25 °C, unless otherwise stated. For induction of gene expression with the Gal4, Gal80^ts^ system, the cultures were carried out at 18°C and then shifted to 29°C prior to dissection. The following lines were obtained from the *Drosophila* Bloomington stock centre at Indiana University: *Or-R (used as wt)*, *w;; Mi(MIC)02608, w;;Df(3R)BSC680, w;;69B-Gal4, w;;UAS-dsRed, w;en-Gal4,UAS-dsRed /CyO, w hsFlp;; FRT 82B tub-GFP. The w;ptc-Gal4-UAS-Gal80^ts^;MKRS/TM6b* was a gift from James Castelli Gair-Hombria, the *w; UAS-wg-GFP* and *w; wg-NRT stocks* were gifts from Jean Paul Vincent. *rn-Gal4*^*13*^ was previously published in St Pierre *et al. 2002*.

### -Transgenic constructs

To generate the *peg-GFP* construct, a fragment containing the *CG17278* 5’ UTR and CDS sequences (devoid of stop codon) was amplified from a whole *Drosophila* embryo cDNA library and cloned into a TOPO-TA gateway pEntry vector, this vector was then recombined with the pPWG vector (UAS-insert-C terminal GFP), obtained from the *Drosophila Genomics Resource Centre at Indiana University*, to obtain *UAS-peg-GFP*. The *UAS-peg-GFP* plasmid was then sent to BESTgene for injection into embryos and generation of transgenic flies.

### -Generation of CRISPR mutants

To generate the *peg* ^−/−^ null mutants we cloned the following guide sequence targeting the CDS of CG17278: GTACCGATCCATTTGCGCTGCGG into the *pU6-BbsI-chiRNA* plasmid, obtained from *addgene*, and following the available protocol (http://FlyCRISPR.molbio.wisc.edu). The following primers were used: *chi-CG17278 ORF guide 1 Fw* CTTCGTACCGATCCATTTGCGCTG and *chi-CG17278 ORF guide 1* Rv AAACCAGCGCAAATGGATCGGTAC. The *pU6-CG17278-chiRNA* plasmid was then sent for injection to BESTgene into the y[1] M{vas-Cas9}ZH2A w[1118] line. Surviving adults were crossed to *w;;Df(3R)BSC680*, the progeny was scored for *peg*−/− like phenotypes, stocks were established from putative *peg* mutants, and *peg* alleles were confirmed by PCR of the locus followed by sequencing.

### -Wing preparations

*For Drosophila* adult wing preparations, the flies were collected in SH media (50% glycerol, 50% ethanol), washed in ethanol and then in dH2O, then wings were then clipped and mounted on a slide with Hoyer’s. The slides were then placed in a hot plate at 65°C for 3-5 hours, with a weight on top of the coverslip to ensure a good flattening of the wings.

### -In situ hybridization

*CG17278* ribo-probe was obtained using the *CG1728 in pEntry* plasmid and its T3 promoter, and the *wg* ribo-probe was obtained from a whole wg cDNA fragment in pBluescript, which was a gift from Joaquin Culi Espigul and Sol Sotillos, it was transcribed using its T3 promoter. Digoxigenin labelling of the ribo-probes was performed with the *Roche DIG RNA labelling mix, Sigma Aldrich*, and the *Promega* T3 RNA polymerase. Wing imaginal discs were dissected in ice-cold PBS and fixed for 20 minutes in 4% paraformaldehyde, and a standard DIG-RNA *in situ* hybridization protocol, as described in Galindo *et al*. 2007, was followed.

**-Antibody stainings:**

For wing imaginal disc immuno-stainings, wandering third instar L3 larvae were dissected by evertion in ice-cold PBS, cleared of digestive system and fat body, fixed for 15 minutes in 4% pfa, and left overnight in methanol at −20°C. The tissues were then washed 3 times with PBS. For wingless stainings, a single wash of 20 minutes with PBS, Tween (0.1%) was performed and the rest of the incubations and washes were carried out with ice cold PBS, BSA (0.2%), and ice cold PBS, respectively, maintianing detergent-free conditions in order to preserve the extracellular signal. Anti-Wg 4D4 (mouse, used at 1:50) and anti-Arm 7A1 (mouse, used 1:50) were obtained from *The Developmental Studies Hybridoma Bank*, at the *University of Iowa*. Anti-Sens (Guinea-Pig, used 1:3000) was a gift from Takashi Koyama. We used the following secondary antibodies: to detect Wg we used anti-mouse biotin, Jackson (1:200), and avidin-Cy5 or avidin-FITC, Jackson, (1:1000) or (1:500) respectively, for other antigens we used anti mouse-Rhodamine, jackson (1:250), and anti guinea-pig-rhodamine, Jackson, (1:250). For nuclear labelling we used DAPI at a final concentration of 300 nM.

Confocal images were acquired on a Zeiss Axio Observer microscope, with an LSM 880 Airyscan module, using a 63x/1.46 oil objective. The signal from Wg, Wg-GFP, Peg-GFP, Arm, and Sens, were always detected using the Airyscan detector. The signal from DAPI and UAS-GFP, or dsRED was detected using the GAasp detectors.

### -Induction of peg and wg-GFP

In order to induce the expression of Wg-GFP and Peg-GFP, *Drosophila* lines of the appropriate genetic backgrounds were crossed to lines carrying *ptc-Gal4-UAS-Gal80*^*ts*^, the progeny was reared at 18°C for aproximately 168h and then shifted to 30°C for 24h. After the shift wandering L3 larvae were collected, dissected in ice cold PBS and fixed in 4% PFA. The rearing and shift times were modified as necessary for the 5h, 8h, 12h, and 48h shifts.

### -Fluorescence Signal intensity measurements

For endogenous Wg, the signal was quantified by counting the total number of Wg particles on 15 successive 2.5 micron ROIs, centred on the D/V boundary. Raw images were thresholded and the particle analyser plugin of imageJ used to obtain the number of particles/ROI. For Induced Wg expression, the fluorescence signal profile was measured within a 20 μm^2^ ROI in the posterior compartment, placed directly adjacent to the *ptc* domain. The Sens fluorescence signal profiles were measured within a region of interest (ROI) of 20 × 60 μm, from the centre of the d/v boundary outwards, on either side of the d/v boundary. For Arm nuclear signal quantification, we calculated the ratio of Arm signal overlapping with DAPI, with that not overlapping it.

### -*in vivo* time lapse imaging

For *in-vivo* time lapse imaging of Wg-GFP diffusion, after induction of expression, we dissected the imaginal discs from wandering L3 larvae of the appropriate genotypes, in ice cold Schneider’s culture medium. The imaginal discs were then transferred to an imaging chamber similar to that used in (*46*); the chamber was constructed by sticking a perforated square of double sided tape on the cover-slip of a cover-slip bottomed culture well (*MatTek Corp*), the orifice of the perforated tape was filled with culture media (Schneider’s media, 2% FBS, 0.2% Penicillin-Streptomycin, 1.25 mg/mL insulin, and 2.5% methyl-cellulose as thickener to reduce drifting of the tissues during imaging). For confocal imaging, 7 section z-stacks were acquired at a rate of a whole stack every 30 seconds, using the LSM 880 Airyscan module of the Zeiss Axio Observer microscope, with a 63x/1.46 oil objective, bleaching was performed after the first stack, by 25 iterations with the 405nm Laser line, at 80% power, in a ROI of 15 × 45nm, placed at an average of 0.5nm from the wg-GFP source. Fluorescence intensity measurements were performed on raw image files, using image J. In order to quantify the fluorescence recovery, we calculated the percentage of the original intensity (pre-bleach) recovered, after bleaching, for each time point, hence the following normalization for each data point: Intensity(t_X_)=(Intensity(t_x_)−Intensity (t_bleach_))*100/ intensity (t_pre-bleach_).

### -FRT-mediated clone generation

In order to obtain *peg*^−/−^ mitotic clones, we generated a *w;; peg* ^*Del 1*^ *FRT82B* recombinant line using the *w hsFlp;; FRT 82B tub-GFP* line as the source of FRT82B. These two lines were crossed, and a 1h 37°C heat shock was induced in the progeny 24h after egg laying.

### -Co-immuno precipitation

60 late third instar larvae per genotype (*da-Gal4*, *UAS-Peg-GFP* or *UAS-GFP*) were homogenized in 400 μl of lysis buffer (20mM Tris-HCl pH 7.4/150mM NaCl/1mM EDTA/ 0.5% NP-40) for 30 min (4°C). Cellular debris was spun at 13 000 rpm for 10 min at 4°C. Supernatant was added to equilibrated GFP-beads (Chromo Tek, NY, USA) and left rotating overnight at 4°C. Beads were washed three times with washing buffer (20mM Tris-HCl pH 7.4/150 mM NaCl/1mM EDTA), and then boiled in Laemmi Loading Buffer (Biorad, CA, USA). Beads were loaded onto 8 or 12% polyacrylamide gel and proteins were separated by SDS-PAGE (Biorad). Detection of specific proteins by Western Blot using a semidry blotting or tetra cell (Biorad). Antibodies used were: mouse anti-GFP 1:2500 (Roche); anti-wg 1:000 (DSHB).

### Quantitative Western blot

For each WB lane, the wing pouches of 10 wing imaginal discs were dissected and homogenised in 20μL of LB2x Buffer (100 mM Tris-HCl pH 6.8, 4% SDS, 0.005% Bromophenol Blue, 20% Glycerol), incubated 5 minutes at 90°C, and centrifuged at 13000 rpm for 5 minutes. The whole lysates were loaded on a Stain Free tgx acrylamide gel (*BioRad*). The proteins were transferred onto a nitrocellulose membrane, with a Trans Blot Turbo Transfer System (*BioRad*) (7 minutes, 1.3A, limited to 25V). Total protein loads were quantified before and after transference, using the Image Lab suite (BioRAD). Wg was detected with anti Wg 4d4 (mouse, DSHB, 1:3000) and anti-mouse-HRP (Jackson, 1:10000), using the ECL select reagent (GE Healthcare) and the Chemidoc MP imager (*BioRad*). Quantification of protein levels were performed against total protein loads for each lane.

### -Quantitative real time reverse transcriptase PCR

Total mRNA was extracted from 30 imaginal discs per genotype, using the RNeasy mini kit (*Quigen*). For each sample, 500ng of mRNA was used for the reverse-transcriptase reaction, using the Quantitect reverse transcriptase kit (*Quiagen*). The qPCRs were performed on a CFX connect thermocycler (*Biorad*) using the Vazyme AceQ SYBR qPCR Master Mix in 20 uL reactions, and using the following primers: Wg fw CCAAGTCGAGGGCAAACAGAA; Wg rv TGGATCGCTGGGTCCATGTA; rp49 fw AGCATACAGGCCCAAGATCG; rp49 rv TGTTGTCGATACCCTTGGGC.

### -Homology searches

For homology searches and phylogenetic analyses we used the same methods as described in (*9*).

### Reaction diffusion model for Wg effective transport

We formalize the effective transport dynamics of Wg with a simple reaction diffusion model,

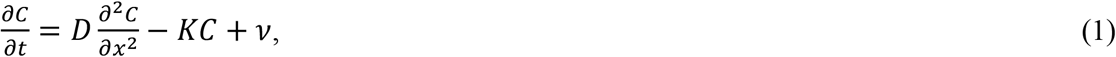

where *C*=*C(x,t)* is the Wg ligand concentration, which is a function of time and space, *D* is the effective diffusion coefficient, *K* is the effective degradation rate and *ν*=*ν(x)* is the ligand source rate, which is spatial dependent. The steady-state solution of Eq. 1 outside of the source (target tissue) for a constant source width and and the length of the tissue in the direction perpendicular to the source being much longer than the decay-length of the gradient is given by,

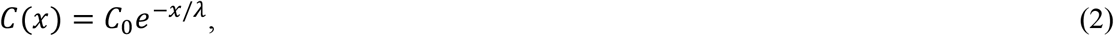

where C_0_ is the amplitude (intensity) of the gradient at the target tissue right next to the source, and λ is the decay length of the gradient, which is a function of D and K in the form,

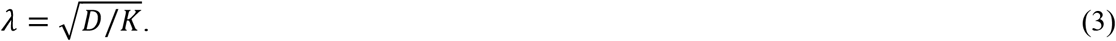

### Analysis of FRAP dynamics

Following the analysis of FRAP dynamics described in [Kicheva et al. 2007], we solved Eq. 1 in space and time. As initial condition at time *t*=*0*, we imposed the steady state profile Eq. 2 outside of the bleached region for *x*<*d* and *x*>*d*+*h*, and *C(x,t = 0)* = *b C_0_ exp(−x/λ)* inside the bleached region for *d*<*x*<*d*+*h*, where *d* is the distance of the ROI from the source, *h* is the ROI width and b is the bleaching depth. Then, the solution of Eq. 1 reads,

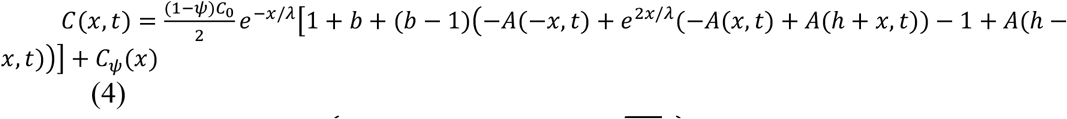

Where 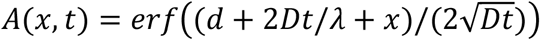 with the error function 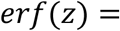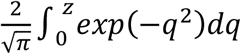, and *C*_*ψ*_(*x*) represents the concentration of immobile molecules that is constant in time, with *C*_*ψ*_(*x*) = *ψC*_*0*_*e*^−*x*/*λ*^ outside of the ROI, and *C*_*ψ*_(*x*) = *bψ C*_*0*_*e*^−*x*/*λ*^ inside of the ROI.

To describe the intensity recovery during FRAP, we calculate the average change in intensity within the ROI in the form 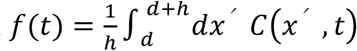. The form of the function *f*(*t*) reads

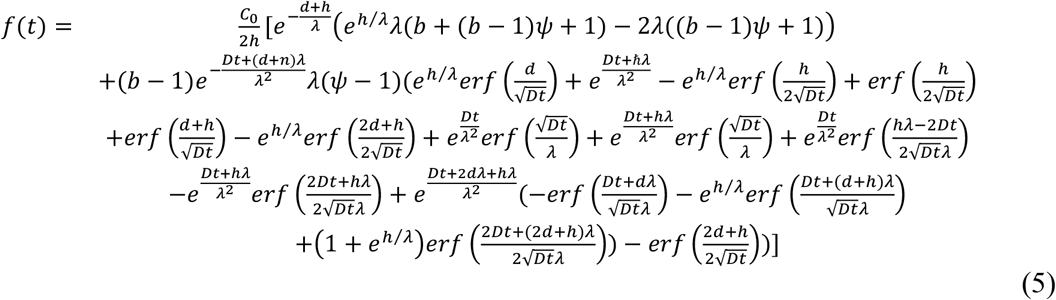

### Determining Wg effective transport dynamics

To find the Wg effective transport dynamics we first fit spatial concentration profiles of endogenous Wg to Eq. 2 in both *wt* and *peg*−/− cases. These profiles were previously normalized with respect to the intensity right next to the source C_0_. This gives values for the decay length λ

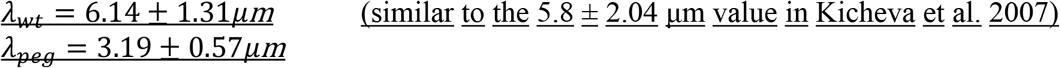

With these values we use Eq.5 to fit the dynamics of FRAP and extract parameters *D*, 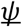 and *b (see table below)*. From the effective diffusion coefficient for both cases

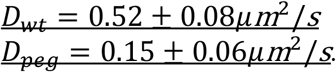

and Eq. 3, we find values for the effective degradation rate

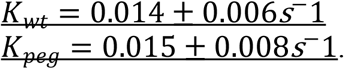

## Supplementary Figures

**Sup. Figure 1.**
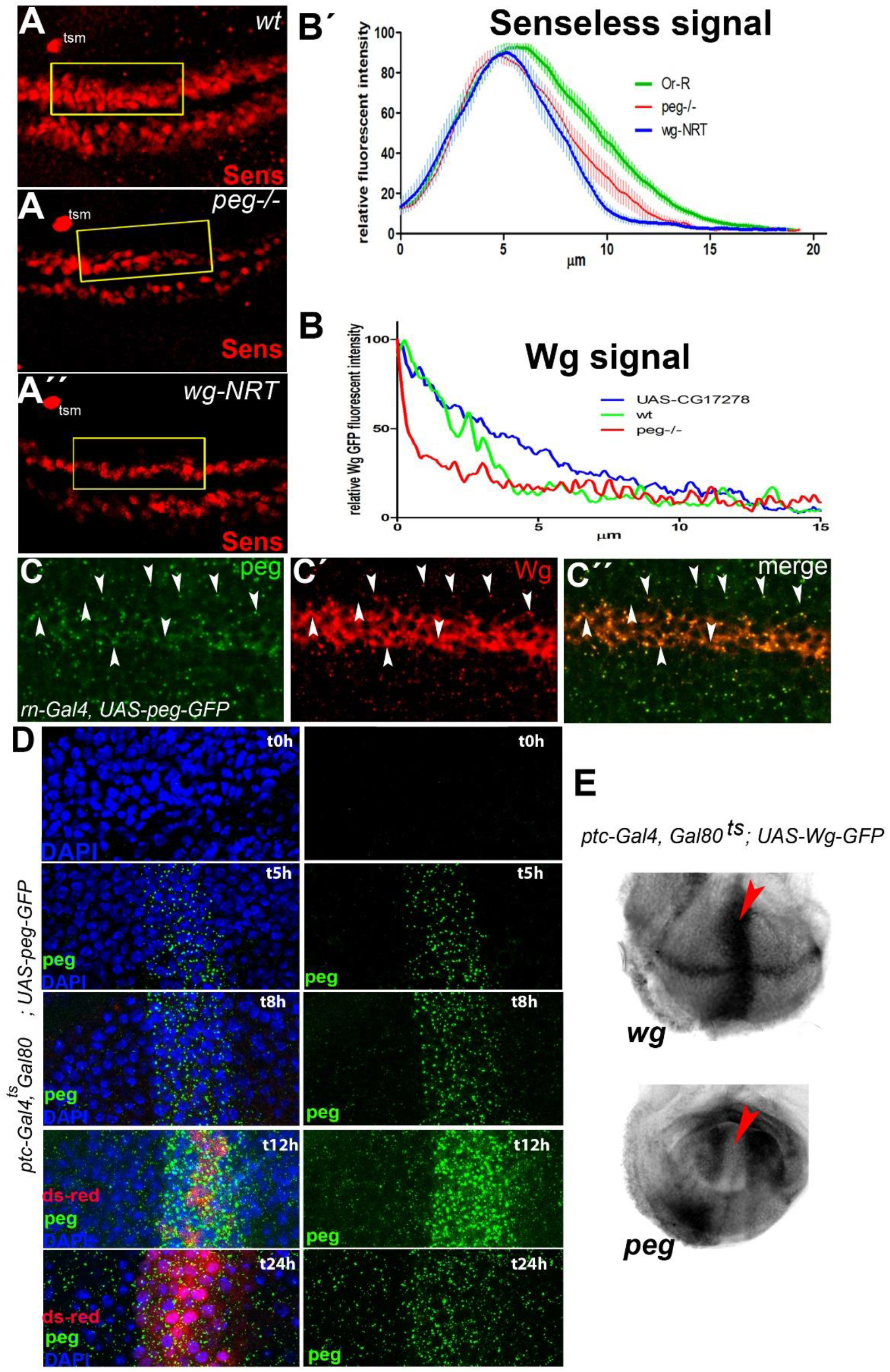
**A-A”.** The expression of *senseless* is significantly reduced in *peg-* mutants and in *wg-NRT* transgenic flies compared to wt. Note that the twin campaniform sensilla (tsm) precursor located some 3-4 cell diameters away from the sens dorsal stripe remains unaffected. **B-B’.** Quantification showing (B) the average Sens signal intensity from panels A-A”, relative to the maximum values. Senseless was quantified in a region of interest, represented by yellow squares in panels A-A”. 0 μm represents the centre of the *wg*-expressing stripe. (B’) quantification of Wg signal, relative to maximum intensities, from (Fig.4 A’-A”) over a region of interest adjacent to the Wg expressing cells. 0 μm represents the boundary of Wg expressing cells. **C.** Endogenous Wg, revealed with the anti-Wg antibody 4D4, co-localizes with Peg-GFP (arrowheads) expressed in the wing pouch with *rn-Gal4(47)*. **D.** Time-line of induction of Peg-GFP expression with *ptc-Gal4-UAS-Gal80*^*ts*^ by temperature shift (18°C to 29°C) prior to dissection shows that Peg-GFP is secreted. The time from induction to dissection is indicated at the right of each set of panels (t). **E.** *In-situ* hybridization showing that ectopic expression of Wg using *ptc-Gal4* leads to a downregulation of *peg* in the *ptc* domain (arrowhead), confirming that *wg* negatively regulates the expression of *peg(24)*.

**Sup. Figure 2.**
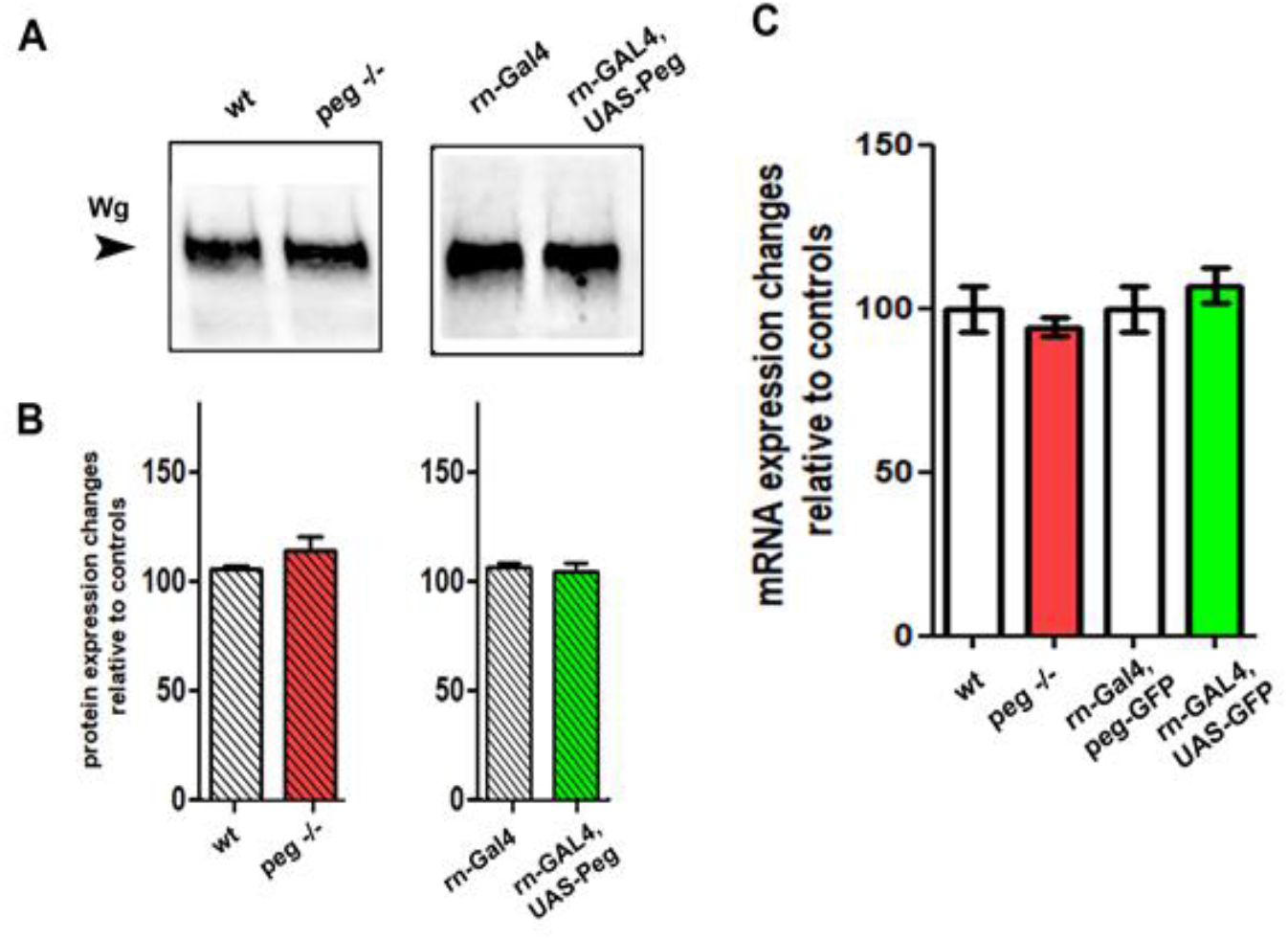
**A.** Western Blots, from wing imaginal disc pouch extracts showing Wg protein levels in *peg*^−/−^ mutants compared to wild-type, or in wing pouches over expressing *peg* with rn-Gal4, compared to controls carrying rn-GAL4 only. Wg levels remain unchanged in either *peg*−/− or *peg* over-expressing imaginal discs, compared to controls. **B.** Quantification of the western blot data from imaginal discs. The Wg protein levels were adjusted against total protein loads, and normalized against the controls. **C.** Quantification of *wg* mRNA levels by qPCR. The values were obtained using the ΔΔQC values of the *wg* amplicon against the *rp49* control and the experimental conditions against the controls. All the values were then normalized to controls. Note that there are no significant *wg* mRNA expression changes between discs mutant for *peg* and wt discs, nor between discs over-expressing *peg* and controls.

**Sup. Figure 3.**
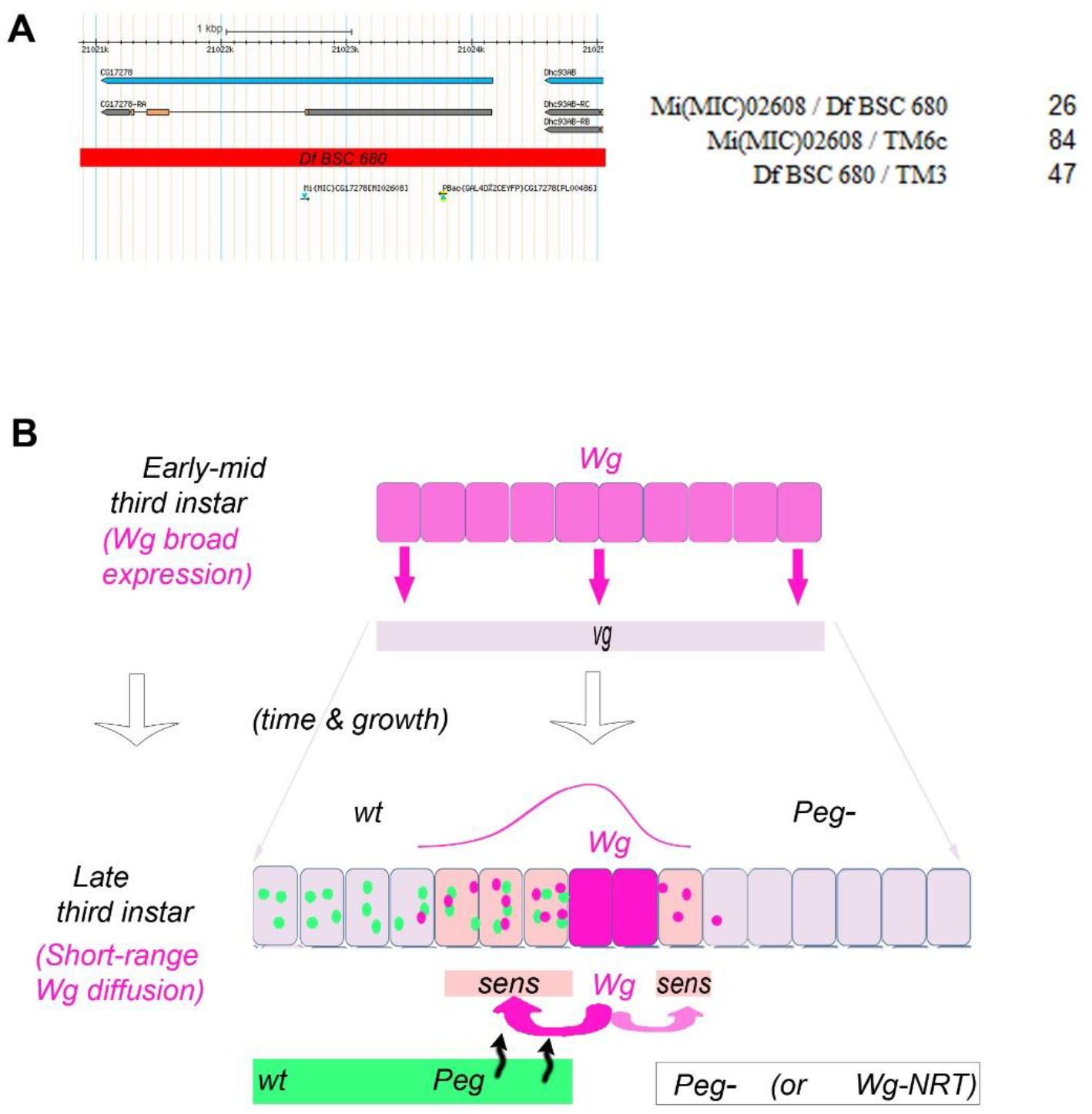
**A.** Genomic locus of *peg*, depicting the sites of the *Mi(MIC)02608* transposon and deficiency *Df(3R)BSC680* used to assess initially the viability of *peg* mutant flies. The table shows the number of progeny of each class obtained after crossing heterozygous flies for both the transposon *Mi(MIC)02608* and deficiency *Df(3R)BSC680*. Note the lower number of trans-heterozygous, *Mi(MIC)02608* / *Df(3R)BSC680* progeny. **B.** Model of *wg* and *peg* function during the development of the fly wing. During Early-mid third instar (aprox. 84 to 96 h. AEL), Wg protein is present in a developing pattern that transitorily reaches the entire developing wing (*20, 34, 48, 49*) and maintains the expression of the *vestigial* gene (*vg*), which is required for the growth of the wing cells. Later, from 96 to 132h AEL, *wg* is expressed only at the presumptive wing margin and is required for its development (see Couso et al. 1994 and references therein). Wg activates the expression of proneural genes such as *sens* (this work) and *ac* at both sides of the *wg*-expressing cells, with an effective range of diffusion of 3-4 cells. Amongst the proneural field, individual cells are selected by a Notch-mediated lateral inhibition process to become the precursors of the wing margin bristles. Chemosensory precursors are determined first by about 120h AEL, and mechanoreceptors later. Complete removal of Wg function from 96h AEL eliminates the entire wing margin (see also (*22*)), whereas later and/or partial removal of Wg reduces the number of chemoreceptors, and then of mechanoreceptors. Peg is required for this process, by being secreted and binding Wg directly it helps Wg effective diffusion up to 3-4 cell away (this work). Loss of Peg reduces the effective range of Wg diffusion to 1-2 cells and produces a mild loss of Wg function phenotype, whereas Wg-NRT reduces it even further (to 1 cell) and produces a stronger, but still not total, loss of Wg function phenotype (Figs. 2 and 5). (see (*50*) and Couso et al. 1994 (*21*) for *wg* wing margin phenotypes). The effective range of Wg diffusion during wing margin development is indicated by the tsm twin sensilla, which is located some 6-8 cells from the *wg*-expressing cells and is not dependant on Wg function (this work; see also Phillips and Whittle 1993).

